# Cortical Dynamics during Contour Integration

**DOI:** 10.1101/2024.05.13.594033

**Authors:** Dongcheng He, Angeline Yang, Daniel R. Coates, Haluk Ogmen, Susana TL Chung

## Abstract

Integrating visual elements into contours is important for object recognition. Previous studies emphasized the role that the primary visual cortex (V1) plays in this process. However, recent evidence suggests that contour integration relies on the coordination of hierarchical substrates of cortical regions through recurrent connections. Many previous studies presented the contour at the same onset-time as the trial, which caused the subsequent neural imaging data to incorporate both visual evocation and contour integration activities, and thus confounding the two. In this study, we varied both the contour onset-time and contour fidelity and used EEG to examine the cortical activities under these conditions. Our results suggest that the temporal N300 represents the grouping and integration of visual elements into contours. Before this signature, we observed interhemispheric connections between lateral frontal and posterior parietal regions that were contingent on the contour location and peaked at around 150ms after contour appearance. Also, the magnitudes of connections between medial frontal and superior parietal regions were dependent on the timing of contour onset and peaked at around 250ms after contour onset. These activities appear to be related to the bottom-up and top-down attentional processing during contour integration, respectively, and shed light on how these processes cooperate dynamically during contour integration.

## 1. Introduction

An important characteristic of natural scenes pertains to the various alignments of edges they contain and the integration of these edges into contours has been thought of as an early step in visual processing to provide an object-level representation. The understanding of contour integration has been guided by the Gestalt principle of good continuation, which highlights that elements of a contour become integrated following certain rules [Wertheimer, 1923], and investigation of the neural correlates encoding these rules has been a longstanding research question. Neurons in early stages of visual processing respond to basic features in the scene (luminance, orientation, spatial frequency, etc.) and their responses are characterized by their receptive fields [Wurtz, 1969; Wilson et al., 1983; Rossi et al., 1996; Ringach et al., 1997; Everson et al., 1998; Hubel & Weisel, 1962]. Although receiving inputs from local, so-called ‘classical receptive fields’, the response properties of these neurons can still be substantially dependent on the stimuli lying outside their classical receptive-fields through lateral feedback and feedforward connections [Gilbert, 1992; Gilbert & Wiesel, 1990; Kapadia, Ito et al., 1995]. These findings lead to the idea that the visual system is able to integrate information from distant cortical areas and provide texture segregation by linking features belonging to the same object.

In support of this idea, previous studies showed that pyramidal neurons in V1 with similar orientation preferences establish intrinsic horizontal connections in spite of non-overlapping receptive fields [Bosking et al., 1997; Schmidt et al. 1997; Stettler, 2002]. Neural models that implement contour integration have also been built around this observation [e.g.: Grossberg & Mingolla, 1985; Ursina et al., 2004; Hansen & Newmann, 2008; Field et al., 1993; Hess & Field, 1999]. These models, based on orientation-selective neurons in V1, consist of neural networks that generate higher outputs when collinear contours or enclosed contours with continuous edges are presented compared to jagged edges. However, contour integration is complex in its variety. Recent studies challenged this theory by showing that jagged contours could be equally salient as collinear contours when corners were placed at the locations with sharp orientation changes [Persike & Meinhardt, 2016; 2017]. These results suggest that contour integration also involves complex features that are represented in different cortical layers other than V1, such as the corners and curves that were found to be encoded in V4 [Jiang et al., 2021]. In another study, Shu-Guang et al. (2017) presented an array of Gabor elements with random orientations and one collinear contour embedded behind a narrow slit. The Gabor array translated horizontally over time and only a vertical band was visible at each frame. In this experiment, subjects showed robust performance in contour detection even when the slit width was equal to the size of a single Gabor element, indicating contour integration at fixed visual fields over time. fMRI data showed engagements of visual areas higher than V1 and posterior parietal areas related to visual memory during the experiments.

In addition to these visual modulations, contour integration has also been found to rely on both top-down and bottom-up attention. For example, Li et al. (2006) reported neural activities from macaque monkeys’ V1 that were dependent on contour saliency, which indicated a bottom-up attention process. As for top-down processing, Chen et al. (2014) found that top-down attention shortened the reaction times of monkeys in a contour detection task. Li et al. (2008) found neural responses that were specific to contour integration in monkeys’ V1 that disappeared when monkeys were under anesthesia. These recent findings suggest that a hierarchical neural substrate could be involved during contour integration, including complex feature extraction, feature binding, memory, attention, etc., and decoding the neural substrates of contour integration has remained an unsolved problem.

Previous neuroimaging studies of contour integration compared behavioral and neural data using stimuli with and without contours, which led to multiple confounding variables. For example, Li et al. (2019) used TMS to disrupt subjects’ V1 and V3B areas at various latencies during a contour integration task. They found that the critical time window, defined as a period during which disruptions to a cortical area caused by TMS resulted in impaired performance in contour detection, could be earlier for V3B than V1. Given the fact that V3B receives information from V1, this finding indicated a recurrent neural path between these two areas during contour integration. However, it is difficult to attribute this recurrent process to specific cognitive processes, since presenting the contour at the same onset-time as the stimulus triggered multiple cortical processes at once with overlapping time windows. Under an ordinary experimental setting with a relatively quiet and dim environment, subjects’ neural activities for visual processing and attentional effort could be simultaneously evoked from being a weak state in response to the onset of the visual stimuli, such as the contour. In another study, Machilsen et al. (2011) examined the effect of contour integration on the Visual Evoked Potential (VEP) by comparing VEP components triggered by contour-absent and contour-embedded Gabor arrays. They found that N1 and P2 components were impacted by the contour. However, these findings did not clearly uncover the neural activities for grouping and integrating visual elements during contour integration, as the visual awareness of a target could induce similar effects on the VEP [Koivisto & Revonsuo, 2010]. One possible way to decouple a signal triggered by a visual target and the large VEP signal triggered by the onset of a stimulus is to place the target with a sufficient latency over periodically changing backgrounds so that these two signals are temporally separated. Subjects’ neural activity will adapt to the background change and thus become relatively small in magnitude compared to the activity triggered by the target.

Therefore, to address these potential confounding variables, in a trial, we separated the contour’s onset from the trial’s onset by presenting a contour-embedded stimulus at various randomized latencies within a sequence of stimuli without the target contour. We also varied the fidelity of the contour to examine how subjects’ neural activities responded to contours with edges having less precise alignment. Furthermore, we recorded EEG signals from subjects during our experiments and investigated the underlying cortical dynamics. First, we conducted an ERP analysis to study how activities in each cortical region changed over time. Next, by considering cortical regions of interest as nodes within a network, we analyzed inter-nodes correlations of EEG data to study how cortical signals change along different cortical pathways during the task. We hypothesized that the contour-triggered cortical activities would temporally synchronize with the onsets of contours, and that signals induced by the integration of visual elements would be dependent on contour fidelity.

## 2. Experiment 1

### 2.1. Methods

#### 2.1.1. Sample Size

To determine the number of subjects and trials to be used in detecting statistically significant ERP components and between-condition differences, Boudewyn et al. (2018) used the Monte Carlo approach to measure the probability of generating significant ERP waveforms and conditioning effects. In their study, one experiment (N=39) used a well-known ERP component, the lateralized readiness potential (LRP), to simulate the ERP with relatively low magnitude (peak magnitude). Results from the Monte Carlo analysis showed that, with 12 subjects each completing 30 trials, the probability of achieving a significant LRP was over 80%. Moreover, using a repeated-measure design, with 12 subjects each completing 45 trials per condition, the probability of achieving a significant between-condition difference with a magnitude above 1.25 μ*V* was over 80%. Based on these results, we recruited 15 subjects in Experiment 1 and 16 subjects in Experiment 2 and ensured that the number of trials per condition were over 45 for each subject. Post hoc results were consistent with these data.

#### 2.1.2. Participants

Sixteen right-handed adults (average age 22.25±2.74 yr; 8 females) with normal or corrected-to-normal vision participated in the study. Fifteen of them participated in the first experiment and all of them participated in the second experiment. These subjects came from the student and researcher populations of the University of California, Berkeley, and included two of the authors: DH and AY. With the exception of the two authors, subjects were unaware of the purpose and hypotheses of the study. All subjects gave written informed consent prior to the commencement of the study. The study was approved by the Institutional Review Board at the University of California, Berkeley.

#### 2.1.3. EEG Apparatus

EEG data were recorded with a Mobita Amplifier (TMSi, Netherlands) from 32 electrodes mounted on a cap according to a 10/20 system. These data were digitized at 1 kHz and referenced online to an electrode placed on the right wrist of subjects during recording.

#### 2.1.4. Experimental Procedures

As illustrated in Figure 1, our stimuli consisted of an array of 27 x 15 white line segments that changed their orientations independently and randomly from 0° to 180°, at a rate of 15 Hz on a black background. The size of each line segment was 1.5 *deg* x 0.15 *deg*, and the separation between centers of any pair of adjacent lines in the same row or column was 2 *deg*. A gray dot (0.5 *deg*) was present at the center of the display for fixation throughout a trial. Each trial comprised 25 frames (1.67 s). During one of the four frames from the 11th (starting at ∼660ms) to the 14th (starting at ∼858ms), the orientations of 12 line-segments became aligned to form an enclosed contour (outline of a box with 3 segments on each side). The center of the contour was at 6 *deg* to the left or right of fixation. Participants were asked to fixate on the dot in the center of the display throughout a trial, and to indicate whether the contour was shown on the left or right of fixation by pressing the corresponding arrow key on the keyboard at the end of a trial. Each block consisted of 120 trials (4 contour onset timings x 2 contour locations x 15 repetitions, presented in a random order), with an inter-trial interval of 1s, during which only the fixation dot was presented. All participants completed three blocks of trials, with an inter-block interval no longer than three minutes. EEG data were collected throughout the experiment. Stimuli were programmed using Matlab (version R2016b, The Mathworks, MA) and Psychtoolbox [Kleiner et al., 2007]. Participants were seated with their heads stabilized on a head and chin rest 70 cm from a 32-inch Display++ LCD monitor (Cambridge Research System Ltd., Rochester, UK) in a dimly lit room. Luminance calibration of the display was performed before data collection, such that the luminance of the line segments, the fixation dot and the background were 148.7, 75.1 (calculation: 74.8) and 0.9 cd/m2, respectively. To synchronize the stimuli displays and EEG signals, digital triggers were sent to the Mobita Amplifier marking temporal onsets of all the trials and the contour-embedded frames. For training purposes, all participants completed one to three practice blocks of 16 trials each, until they had fewer than two response errors within a block. All participants passed the training session within three blocks. EEG data were not recorded during training and the psychophysical data obtained during training were not included for analysis in this paper. The entire experiment (including training) took approximately 30 minutes to complete.

**Fig. 1.**
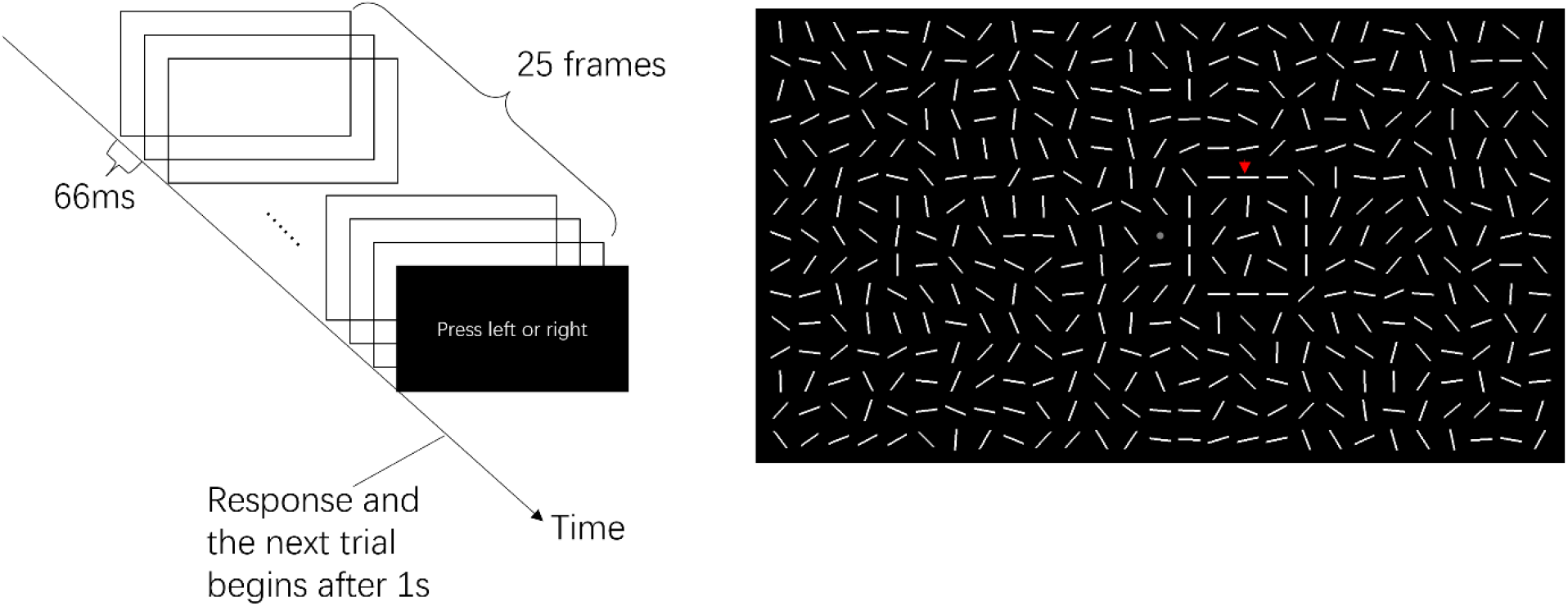
Illustration of the experimental paradigm and an exemplar frame presenting the contour on the right side to the fixation dot, as marked by a red arrow.

#### 2.1.5. ERP Data Preprocessing

Offline data analysis was performed using MNE, a Python package for neurophysiology data analysis and visualization [Gramfort et al., 2013]. EEG data were first band-pass filtered between 1 and 50 Hz. Each trial was then extracted as an epoch starting at 100ms prior to the trial onset and ending at 1500ms after the trial onset. Epochs associated with incorrect trials were removed. Then, signals were re-referenced to the average value across all electrodes. Artifact rejection was performed based on one author’s observation and epochs subject to contamination due to eye blinks, eye movements, myogenic activities, or electrode contacts at any electrode were excluded from subsequent analyses. Finally, each epoch was baseline-corrected against the mean voltage of an interval from 100ms prior to trial onset to 600ms after trial onset.

#### 2.1.6. Temporal Data Analysis of ERP

We selected epochs from six electrodes (FZ, P3, P4, T7, T8, OZ) to identify the temporal dynamics of EEG data localized to the regions of interest. For the electrodes placed on the midline (FZ and OZ), the remaining epochs after pre-processing were averaged for each contour-onset condition. As for other electrodes placed on posterior parietal and temporal regions, data were averaged based on both the onset of contour and its lateralization. For each contour onset condition, we defined contralateral channels, PC and TC, as averaged epochs over trials in which contours were presented on the contralateral side to the electrodes. For example, PC represented averaged epochs from electrode P3 over the trials when the contour was presented to the right of fixation dot and electrode P4 over the trials when the contour was presented to the left of fixation dot. Similarly, we also defined ipsilateral channels, PI and TI, to represent averaged epochs on these four electrodes placed on posterior parietal and temporal regions over trials in which contours appeared on the same lateralization of the display as the electrodes of the hemisphere. In addition, to examine the effect due to the timing of contour, we extracted the averaged epochs of each subject for all the channels and synchronized these data by the onset of the contour.

#### 2.1.7. Inter-ROI Correlations of ERP Activity

As depicted in Figure 2, based on preliminary results, we defined ten regions of interest (ROIs) enclosed by electrodes, including prefrontal (PF), medial frontal (MF), left lateral frontal (lLF), right lateral frontal (rLF), superior parietal (SP), left temporal (lT), right temporal (rT), left posterior parietal (lPP), right posterior parietal (rPP), and occipital (O). Based on these ROIs, we defined a simplex, a network concept, as two interconnected ROIs, in which the strength of their connection was measured by the correlations of neural activities (EEG signals) between these ROIs. To capture the temporal evolution of each simplex, we paired all the electrodes within one ROI with each electrode within the other ROI. For each electrode pair, moving time correlation was performed on the ERP data starting at the trial onset and ending at 1500ms afterwards using a Pearson correlation method with a time segment of 100ms, a time interval of 1ms, and 0 lag. The inter-ROI correlation of a simplex was defined as the average correlation for all the electrode pairs connecting these two ROIs. These procedures were repeated for each subject, contour timing condition, and simplex.

**Fig. 2.**
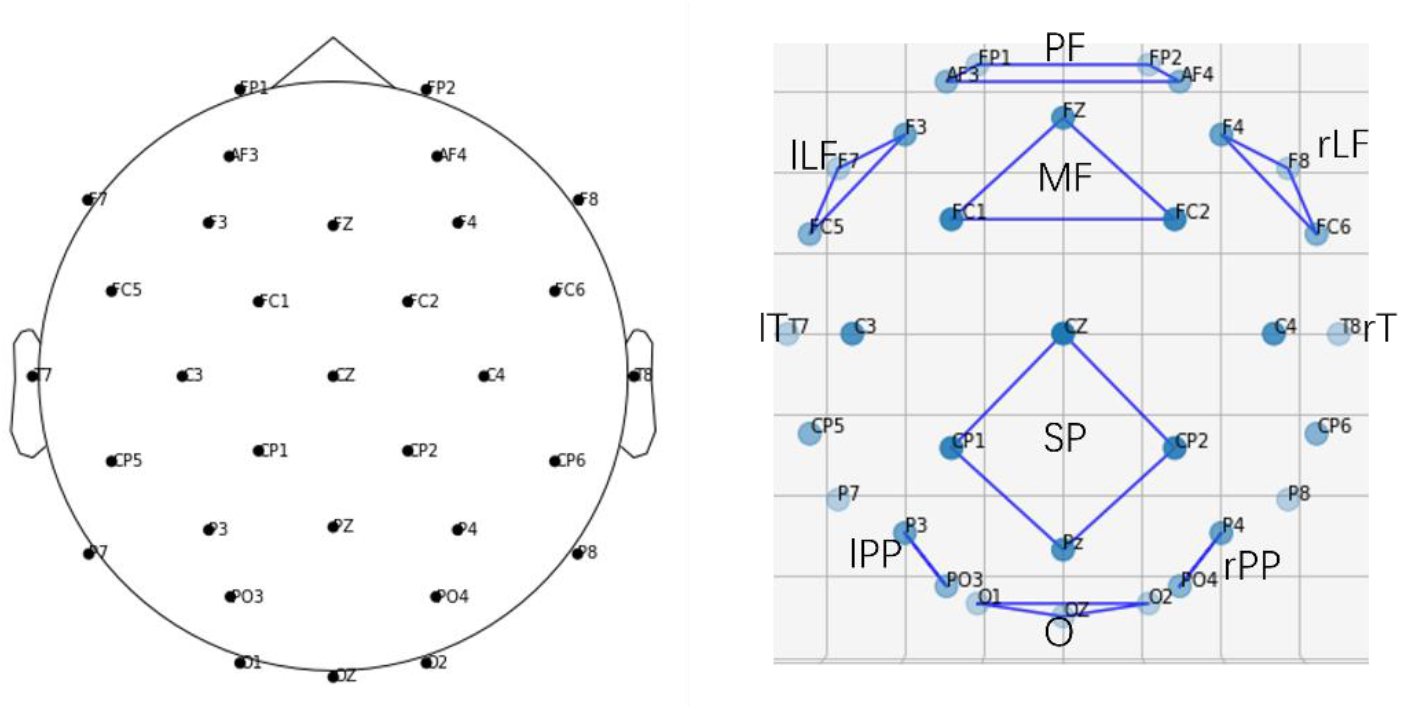
Electrode locations relative to the scalp and regions of interest as defined by enclosure of electrodes. Prefrontal (PF): FP1, FP2, AF3, and AF4; medial frontal (MF): FZ, FC1, and FC2; left lateral frontal (lLF): F3, F7, and FC5; right lateral frontal (rLF): F4, F8, and FC6; superior parietal (SP): CZ, CP1, CP2, and PZ; left temporal (lT): T7; right temporal (rT): T8; left posterior parietal (lPP): P3 and PO3; right posterior parietal (rPP): P4 and PO4; occipital (O): O1, O2, and OZ.

### 2.2. Results

The mean behavioral performance across subjects in this experiment was 98.83%±2.08% correct. After the onset of the contour-embedded frame, we identified that frontal P200, occipital N200 and P400, posterior parietal P400 and contralateral N200, as well as temporal N300 were synchronized with the onset timing of the contour (Figure 3). Information about the EEG channels covering these regions are illustrated in the Methods section. T-tests showed that the peak values of all these ERP components were significantly larger than zero, with results shown in Table 1. For each of the channels (FZ, PI, PC, TI, TC, OZ), we aligned the ERPs by the onsets of the contour embedded frame, as shown in Figure 4, and ran a repeated-measures ANOVA with the onset timing as the main factor on the ERP magnitudes of each time sample. After multiple-testing correction on a basis of 20ms, significant effects of contour onset timing were detected on signals from PC during the period from 521 to 533ms, and signals from PI during the period from 376 to 386ms. These effects detected in the posterior channels persisted no longer than 12ms, other than which no significant effect of contour onset timing was found in any channel and latency. In spite of the relatively weak temporal effects on posterior signals, these results suggest that the ERPs we identified above were independent of temporal factors such as subjects’ expectations about the stimulus onset and the sequential order of the stimulus.

**Fig. 3.**
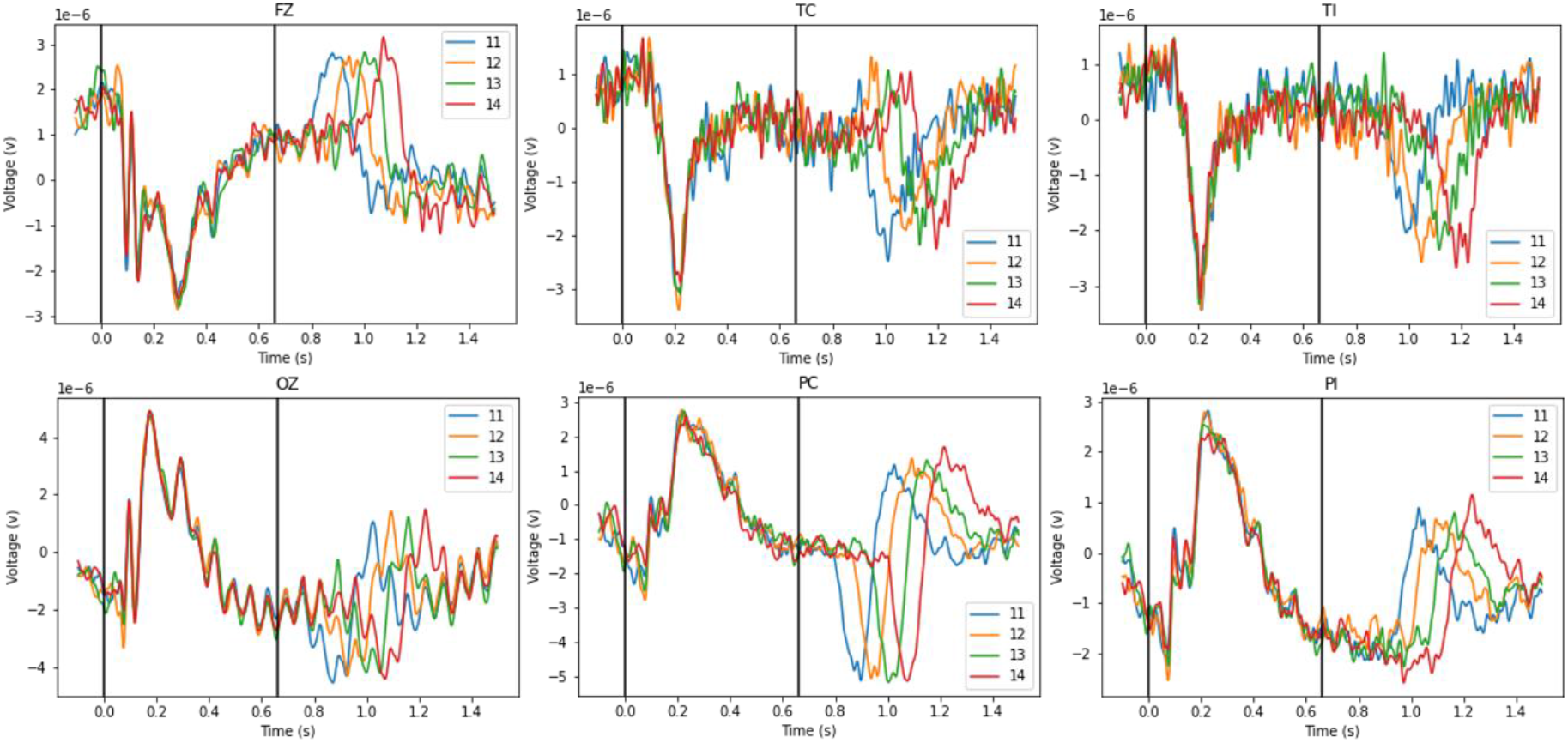
Average ERPs across all trials and subjects in six channels. FZ and OZ are labeled by original electrodes, whereas other four channels are reproduced based on the laterization effect. Channel TC and TI are reproduced from electrodes T7 and T8, depending on whether the contour appears on the contralateral (TC) or ipsilateral (TI) side of the electrode. Similarly, PC and PI are reproduced from electrodes P3 and P4. Different colors represent different temporal orders of the contour, as labeled by the ordinal number of the contour embedded frame. One vertical line marks 0ms, the onset of trial. A second vertical line marks 660ms, the onset of 11^th^ frame.

**Fig. 4.**
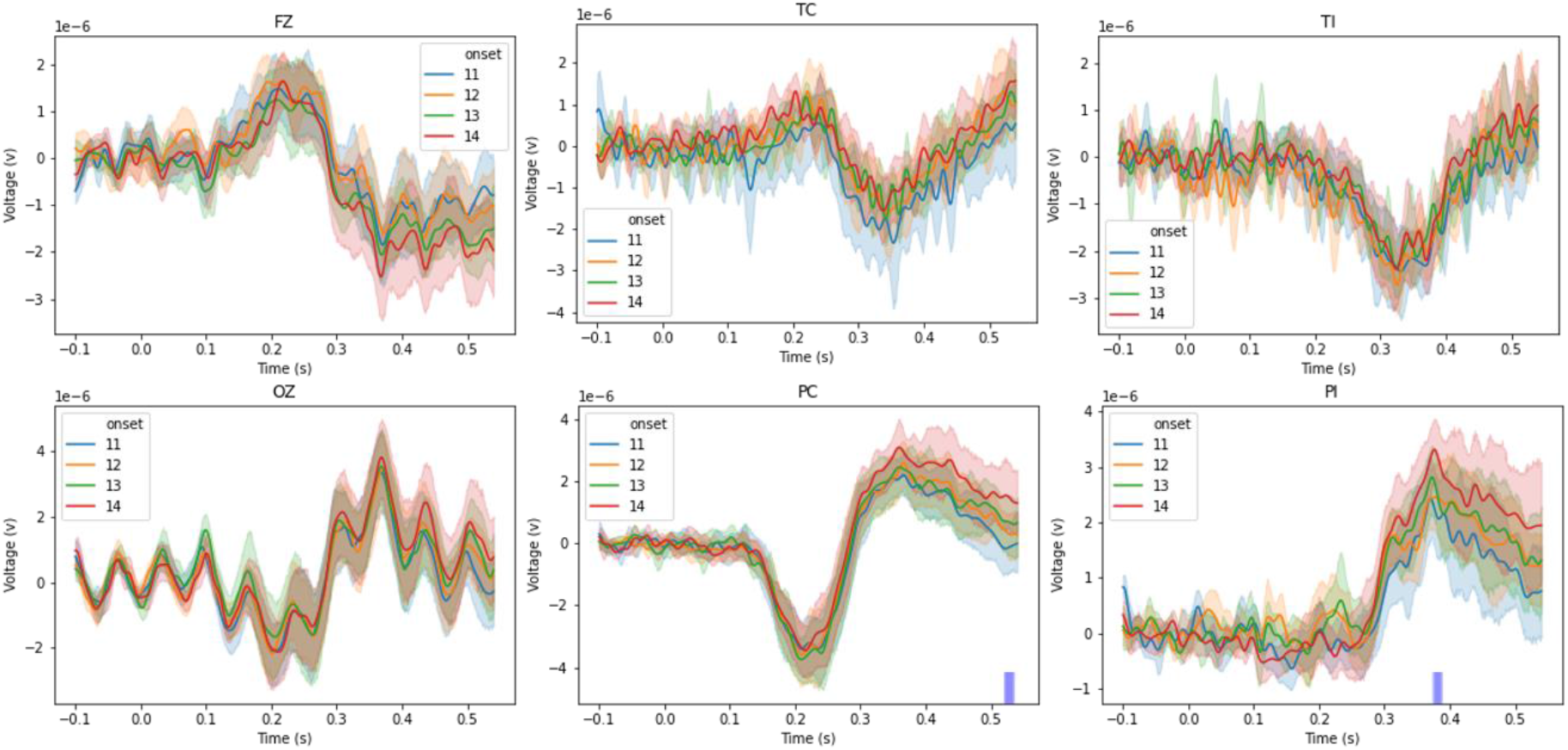
Average ERPs aligned by the onset of the contour embedded frame, as represented by 0ms. Shadings indicate the 95% of confidence interval across all subjects. Channel labels and color legends are same as Fig. 3. The purple bars lying on the X axis mark the latency period when RM-ANOVA detected a significant effect from onset conditions on ERP amplitudes after being corrected by an interval of 20ms with a critical value of 0.05.

**Table 1:** ERP Components and Results of T-test Comparing Peak Values to Zero.

| ERP Component | Peak Latency (ms) | \|Peak Value\| ($\mu V$) | T-Test (df=14) |
| --- | --- | --- | --- |
| FZ P200 | 214 | $1.62 \pm 1$ | $T=5.82, p<0.01$ |
| OZ N200 | 202 | $2.18 \pm 1.84$ | $T=4.26, p<0.01$ |
| OZ P400 | 368 | $3.43 \pm 2.03$ | $T=6.08, p<0.01$ |
| TC N300 | 341 | $1.68 \pm 1.23$ | $T=4.92, p<0.01$ |
| TI N300 | 325 | $2.32 \pm 1.01$ | $T=8.25, p<0.01$ |
| PC N200 | 211 | $3.53 \pm 1.5$ | $T=8.48, p<0.01$ |
| PC P400 | 363 | $2.56 \pm 1.16$ | $T=7.94, p<0.01$ |
| PI P400 | 372 | $2.73 \pm 1.04$ | $T=9.46, p<0.01$ |

We also observed strong periodical oscillations in the signals from the electrode OZ, as shown in Figures 3 and 4. To examine the cause of these oscillations, we ran a Fast Fourier Analysis on each subject’s signals from OZ between 0 and 1500ms after contour onsets. The results, shown in Figure 5, indicated that these oscillations were dominated by the 15Hz component, which was equivalent to the rate of orientation changes of our stimuli. Additionally, peaks at its second (30Hz) and third (45Hz) harmonics are also observable from Figure 5. Therefore, we attribute these oscillations to the visual responses directly induced by the temporal periodicity within the stimulus.

**Fig. 5.**
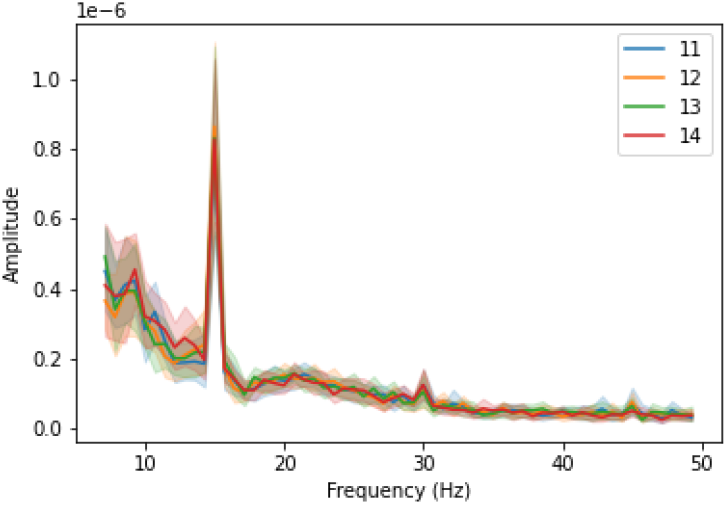
Results of Fast Fourier Transform Analysis. Shadings indicate the 95% confidence intervals across participants. The dominant peak is at 15Hz.

Among all the simplexes, we identified six: (1) lLF-rPP, (2) rLF-lPP, (3) O-lPP-rPP, (4) O-PF-MF, (5) SP-MF, and (6) lT-rT, whose activities synchronized with the onset of the contour and showed robust stronger or weaker inter-ROI correlations between 100ms and 300ms after the onset of the contour. These results are shown in Figures 6 and 7, where we aligned these inter-ROI correlations for each simplex by the onsets of trial and contour, respectively. Using the data from contour onsets to 500ms after contour onset, we ran a repeated-measures ANOVA with the onset timing as the main factor on the ERP magnitudes of each temporal sample. After multiple-testing correction on a basis of 20ms intervals, only the correlations between SP and MF showed a significant effect of contour timing and its duration was over the period from 220 to 250ms (peaked at 225ms: F(3,39)=4.09, p=0.01). The amplitude of the SP-MF connection was reduced after contour appearance, with lower mean peak values for later contour onsets. This suggests that the dynamics of the SP-MF pathway were contingent on the event uncertainty, as subjects’ uncertainty about contours’ appearance reduced with respect to the timing. When subjects first observed a contour in an early frame, there was still a probability that this observation was a false-alarm and that the actual contour could appear later. However, such probability was zero if subjects first observed the contour at the last possible frame (14th frame).

**Fig. 6.**
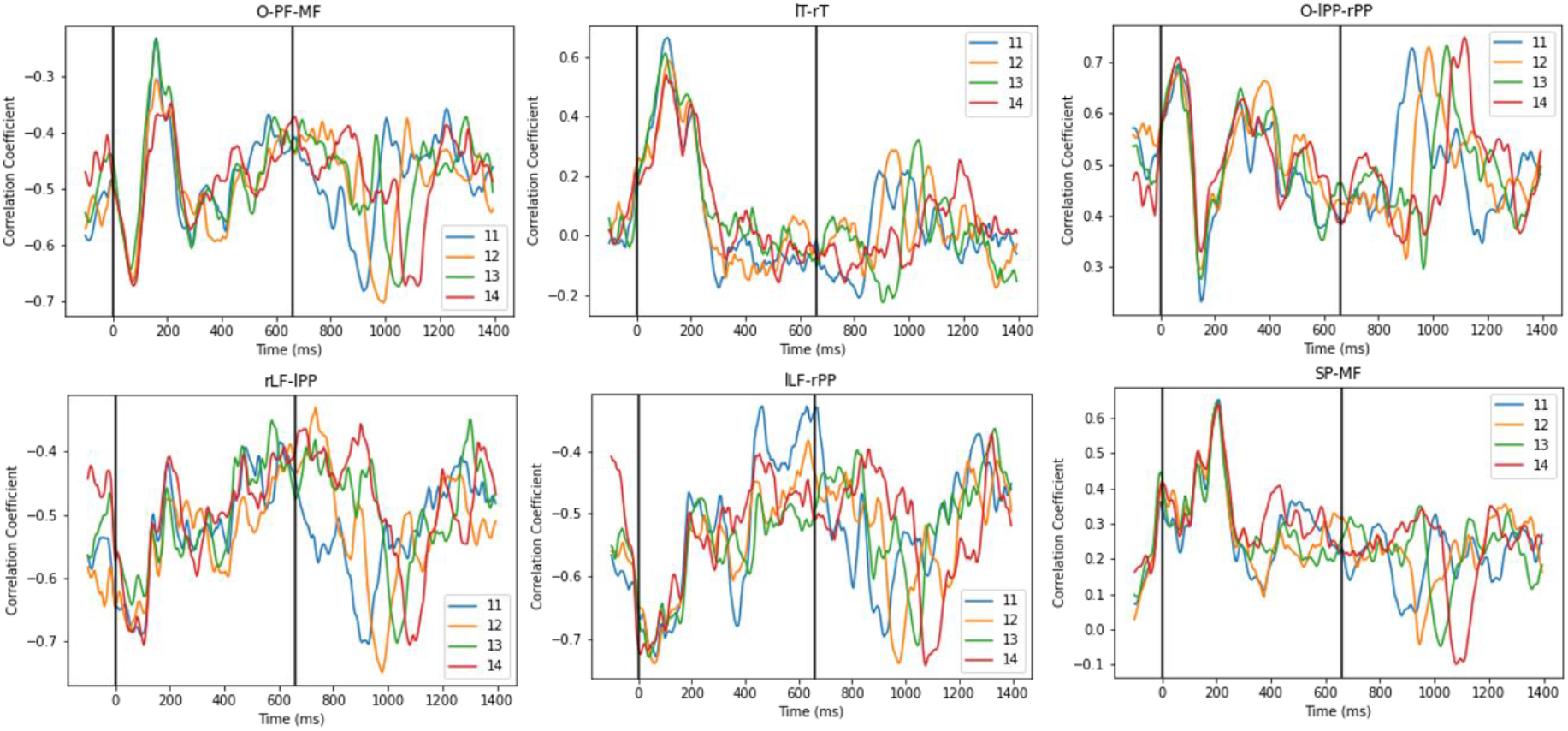
Average inter-ROI correlations across all trials and subjects. One vertical line marks 0ms, the onset of trial. A second vertical line marks 660ms, the onset of 11^th^ frame.

**Fig. 7.**
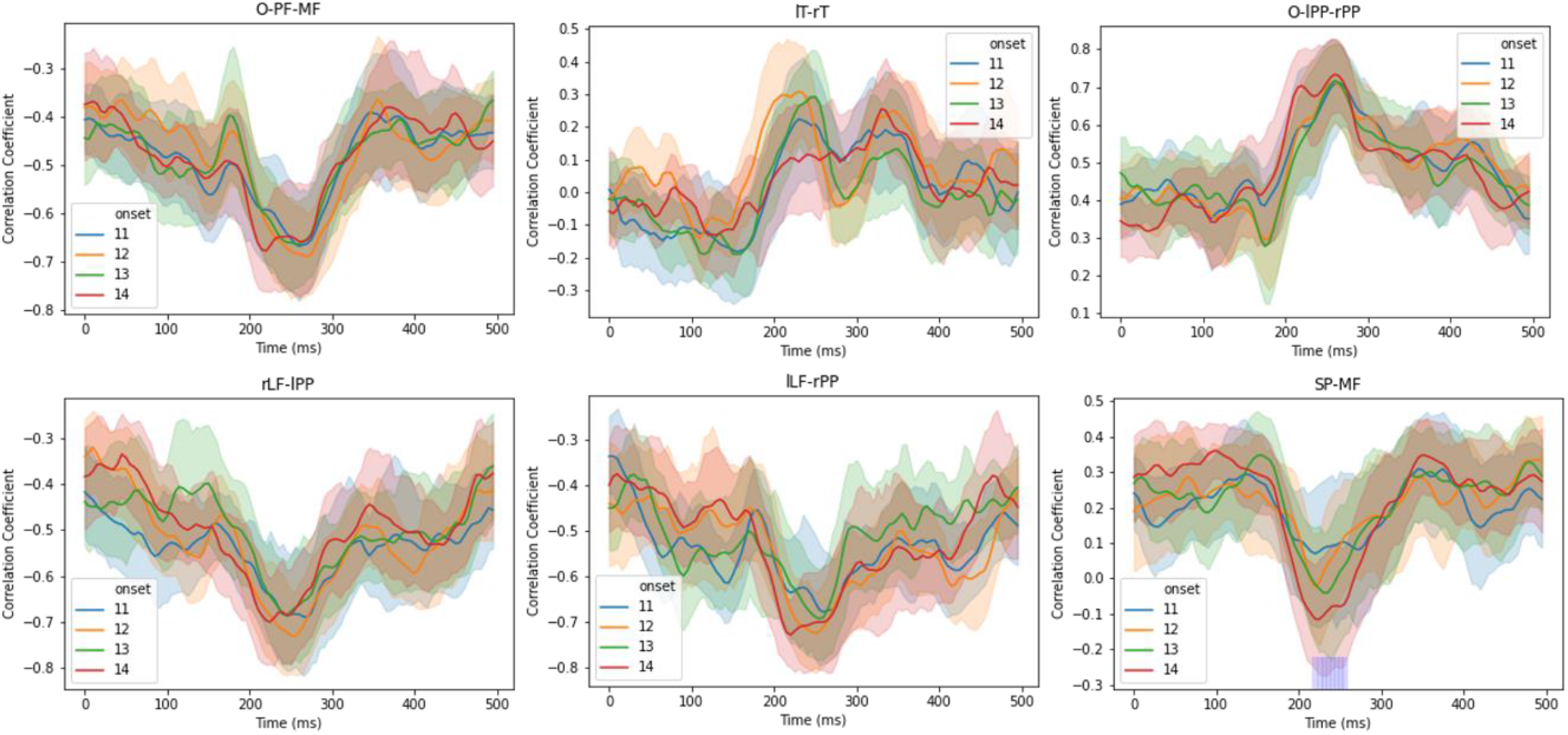
Average inter-ROI correlations aligned by the onset of the contour embedded frame, as represented by 0ms. Shadings indicate the 95% of confidence interval across all subjects. Color legends represent the ordinal number of contour embedded frame. The purple squares along the X-axis in the bottom right panel mark the latency period when RM-ANOVA detected a significant effect from contour onsets on inter-ROI correlations after being corrected by an interval of 20ms with a critical value of 0.05. Notations for ROIs can be found in Fig. 6.

To compare these simplex dynamics through time, we normalized these data by taking the absolute values of the correlation coefficients and baseline-corrected them to the values at 100ms prior to contour onset, and then averaged these values across subjects and contour-onset conditions (Figure 8). We categorized correlation coefficients from lLF-rPP and rLF-lPP into a contralateral class (CL) if the stimulus was presented contralateral to the corresponding side of the posterior region, and an ipsilateral class (IL) when the stimulus and the posterior region were ipsilateral. We found both pathways had a contralateral effect that synchronized to the contour onset, with a peak correlation at around 150ms. Next, around 200 to 300ms, we observed correlations above baseline among the occipital, frontal, and lateral posterior areas, increased bilateral temporal areas correlation, as well as reduced correlations between superior-inferior parietal and median frontal areas. All correlations returned to baseline levels within 500 ms after contour appearance. Peak values of the normalized Inter-ROI Correlations are shown in Table 2.

**Fig. 8.**
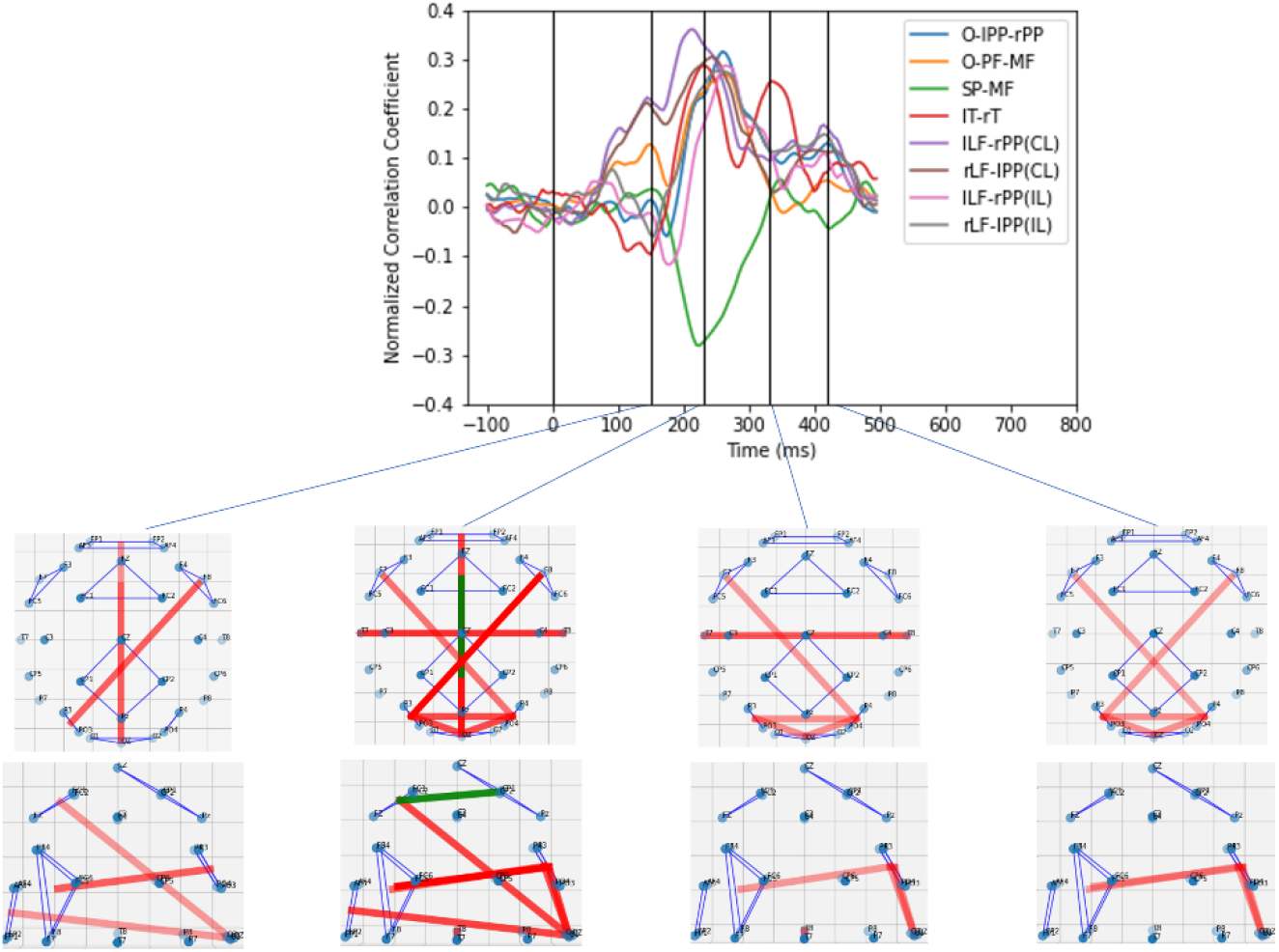
Normalized inter-ROI correlations aligned by the onset of the contour embedded frame, as represented by 0ms. Color legends represent simplexes. The cortical pathways from four instants (150ms, 230ms, 330ms, and 420ms) for presenting the contour on the right side are visualized in the topographies, with both top views (first row) and side views (second row). Red edges indicate above-baseline activities and green edges indicate below-baseline activities. The transparency of edges is contingent on the absolute values of corresponding normalized correlation coefficients, of which values below 0.1 are not visualized.

**Table 2:** Normalized Inter-ROI Correlations and Results of T-test Comparing Peak Values and Zero.

| Simplex | Peak Latency(ms) | \|Peak Value\| | T-Test (df=14) |
| --- | --- | --- | --- |
| O-IPP-rPP | 260 | 0.32±0.2 | T=5.51, p<0.01 |
| O-PF-MF | 260 | 0.27±0.18 | T=5.62, p<0.01 |
| SP-MF | 225 | 0.28±0.3 | T=3.52, p<0.01 |
| IT-rT | 230 | 0.29±0.25 | T=4.37, p<0.01 |
| ILF-rPP [CL-IL] | 195 | 0.4±0.38 | T=3.94, p<0.01 |
| rLF-IPP [CL-IL] | 150 | 0.26±0.15 | T=6.4, p<0.01 |
| ILF-rPP [(CL+IL)/2] | 255 | 0.27±0.19 | T=5.51, p<0.01 |
| rLF-IPP [(CL+IL)/2] | 245 | 0.29±0.2 | T=5.41, p<0.01 |

### 2.3. Conclusions

In this experiment, we separated the EEG signals triggered by the contour’s appearance from those triggered by the onset of the stimulus display. As shown in Figures 3 and 6, cortical dynamics exhibited different patterns after the onset of stimulus display and the onset of contour. By varying the timing of contours’ onsets, we identified multiple signatures that synchronized with the contours’ onsets in the temporal domain, which were thus considered to be related to contour integration. For the ERPs, we identified frontal P200, temporal N300, occipital N200, posterior parietal contralateral N200, and posterior P400. For the inter-ROI correlations, we found strong correlations between the posterior and frontal regions after contours were presented. Among all these, we observed two interesting phenomena. First, our data showed interhemispheric correlations between the lateral frontal and posterior parietal regions with peak magnitudes at around 150ms after the contours’ onsets, whose lateralization was dependent on contours’ location. Second, we found the correlations between the median frontal and superior parietal regions to be dependent on the temporal order of the contour, with peak magnitudes at around 250ms after contour onset. These findings implied how the bottom-up and top-down attentional controls were deployed cooperatively and we will discuss this in detail in the Discussion section.

The results of this experiment suggest the involvement of cortical activities across broad regions over the entire cortex during the contour integration process. To answer the question of which signatures directly represent the grouping and integration of visual elements into contours, in the next experiment, we varied the fidelity of contour and examined how the ERP signatures we identified would be impacted. We hypothesized that the magnitudes of ERP signals induced by the perceptual grouping and integration of line segments of the contour should depend on contour fidelity. On the other hand, activities induced by other cognitive functions should be independent of contour fidelity.

## 3. Experiment 2

### 3.1. Methods

#### 3.1.1. Experimental Procedures

In this experiment, we varied the fidelity of contours by adding random noise in the form of orientation jitter to the line segments that formed the contour stimuli. As a measure of contour fidelity, we defined the degree of orientation jitter by the range of angular disparities that applied to all the line segments in a contour. As an example, in the case of a jitter of 30°, a sequence of twelve angular disparities lying in the range of [-30°, +30°] was randomly generated to be added to the original orientations of the 12 line segments that formed the enclosed contour, as depicted in Figure 9.

**Fig. 9.**
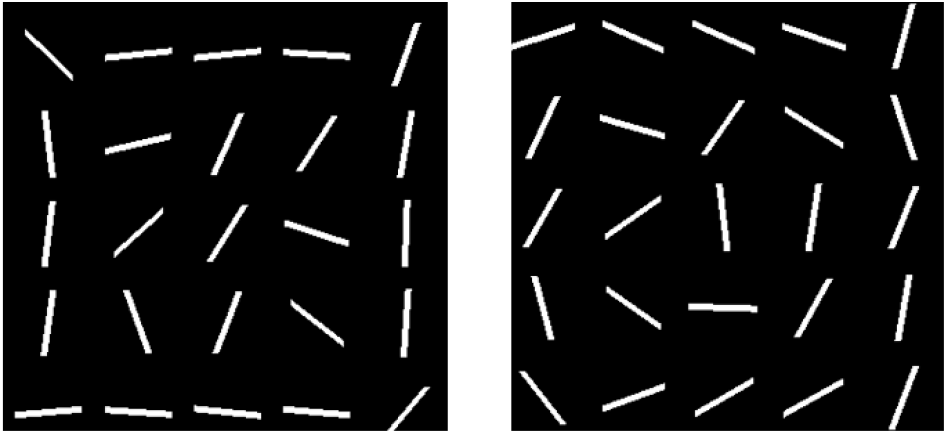
Examples of contours with two different fidelities. The variation degrees of the left and right figures are 10° and 30°, respectively.

This experiment consisted of two sessions. In the first session, we measured each subject’s psychophysical performance (EEG was not recorded) in response to the amount of orientation jitter of the contour elements. The purpose of this session was to determine the threshold orientation-jitter for each subject to be used in the second session of the experiment in which EEG recording would be performed. Stimulus configuration and experimental procedures were similar to those of Experiment 1, with two exceptions: (1) each trial comprised 20 frames (1.32 s) and the contour only appeared in the 11th frame (started at ∼660ms); and (2) contours were presented in five levels of orientation jitter: 0°, 15°, 30°, 45°, and 60°. This session contained only one single block of 200 trials (5 orientation jitters x 2 contour locations x 20 repetitions) in a random order. For each subject, a psychometric function relating their performance (percent-correct of identifying the right/left location of the contour) with the orientation jitter of the contour was created by fitting these data with a Weibull distribution function, assuming the lapse rate was 0, reparametrized by a chance level at 50% [May & Solomon, 2013]. From the fitted psychometric function, we determined the threshold jitter that corresponded to a performance accuracy of 75%. This jitter value was used in the second session in which EEG data were collected while subjects identified the location of the contour. Experimental procedures were similar to those in the first session. For each subject, we tested three contour fidelities: full, partial, and null, which corresponded to three orientation-jitter conditions: 0°, *t*ℎr*es*ℎ*old jitte*r, 180°, respectively. This session consisted of two blocks of 90 trials each (3 orientation jitters x 2 contour locations x 15 repetitions) in a random order. On average, subjects completed both sessions in an hour.

#### 3.1.2. ERP Data Pre-processing and Analysis

Unless otherwise stated, pre-processing and analysis of ERP data were similar to those of Experiment 1. EEG data were extracted into epochs starting at 500ms prior to the onset of the contour and ending at 700ms after the onset. Epochs associated with incorrect trials were removed for orientation jitter of 0° and the threshold jitter value (not for 180°). Each epoch was baseline-corrected against the mean voltage of an interval from 200ms prior to contour onset to 200ms after that. To facilitate comparison of ERP signals across the varying contour fidelities (other than 180°), we normalized the amplitude of each ERP signal by taking into account subjects’ performance and noise level. This adjustment aimed to boost the signal-to-noise ratio and reveal the magnitude of the effect that was unaffected by the trials in which subjects were correct based simply on guessing, as explained by the following formulae.

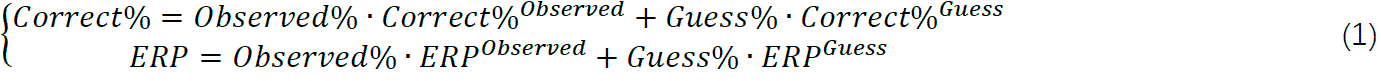

where “Observed” indicates the trials in which subjects genuinely observed the location of the contour, and “Guess” indicates the trials in which subjects were correct based on guessing. To uncover and compare *E*RP^*Obse*r*ved*^, we used the measured signals for the orientation jitter of 180°, *E*RP^180°^, to represent *E*RP^*Guess*^, and adjusted the original *E*RP using the following assumptions and equations:

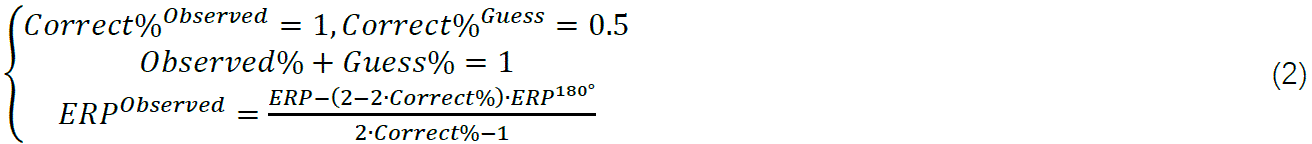

Finally, to reject the high frequency oscillations and random noise and to compare the ERPs among different fidelity conditions, we used an equal-weighted filter with a length of 200ms to smooth all the ERPs by convolution.

### 3.2. Results

Psychometric functions obtained from the first session of all subjects are plotted in Figure 10. The threshold orientation-jitter for the 16 subjects ranged between 18 deg and 45.75 deg (mean value: 32±7.8 deg). Subjects’ mean performance in the second session for the three orientation jitter conditions were 98.23%±3.13% for the 0 deg (“full”) condition, 80.21%±5.47% for the threshold-jitter (“partial”) condition, and 47.29±5.9% for the 180 deg (“null”) condition.

**Fig. 10.**
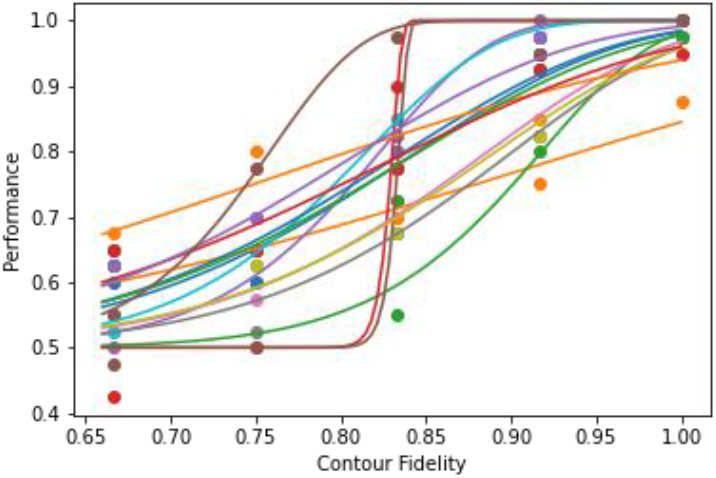
Psychometric functions of subjects. The horizontal axis, labeled as Contour fidelity, indicates the fidelity level measured by 1-variation degree/180. Each color represents a subject.

We merged the ERPs from electrodes T7 and T8, as N300 from these electrodes were unaffected by the contour’s lateralization, as revealed in Figure 3 and 4. ERPs were then aligned by the onsets of the contour-embedded frame and are visualized in Figure 11 and 12, where we showed ERPs before and after the performance-based adjustment, respectively. For all these channels, no salient ERPs were observed in the null condition. Furthermore, to examine the effect of contour fidelity, we ran a repeated-measures ANOVA with the fidelity condition (full versus partial) as the main factor on the ERP magnitudes of each time sample. After multiple-testing correction on a basis of 20ms intervals, in Figure 12, we observed a significant effect of fidelity for the P400 components in channels PI (at mean peak latency 422ms: F(1,15)=17.43, p<0.01) and OZ (at mean peak latency 382ms: F(1,15)=25.34, p<0.01), and another significant effect for the temporal N300 (at mean peak latency 366ms: F(1,15)=13.03, p<0.01). These results show that both temporal N300 and late posterior P400 are sensitive to the strength of grouping stimulated by the contours’ quality in shape, whereas other ERPs are not directly related to such factors.

**Fig. 11.**
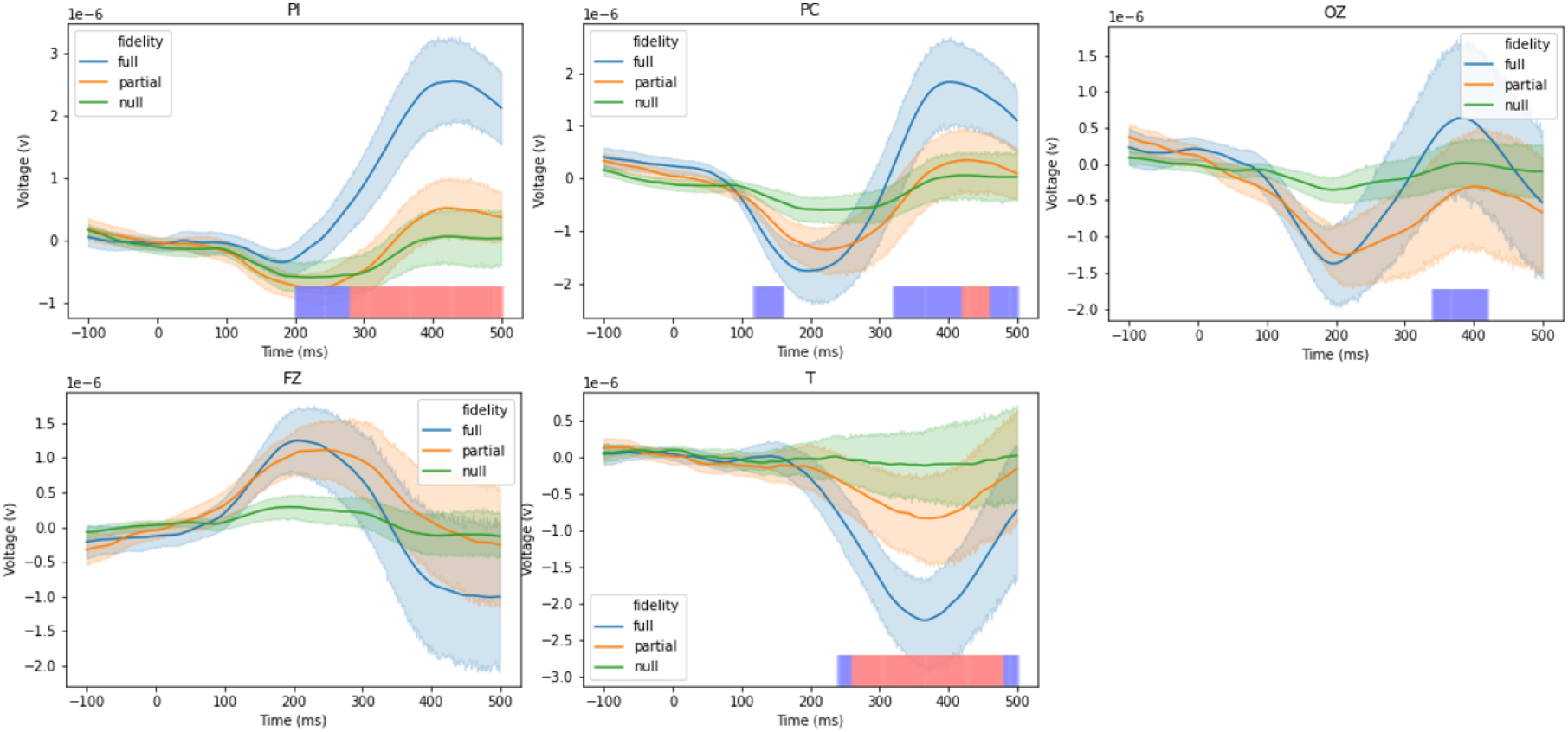
Average ERPs before performance-based adjustment. These signals are aligned by the onset of the contour embedded frame, as represented by 0ms. Shadings indicate the 95% of confidence interval across all subjects. Different colors represent three contour fidelity conditions. The purple and pink squares lying on the X-axis mark the latency period when RM-ANOVA detected a significant effect from fidelity conditions (full vs partial) on ERP amplitudes after being corrected by an interval of 20ms with a critical value of 0.05 (purple) and 0.01 (pink).

**Fig. 12.**
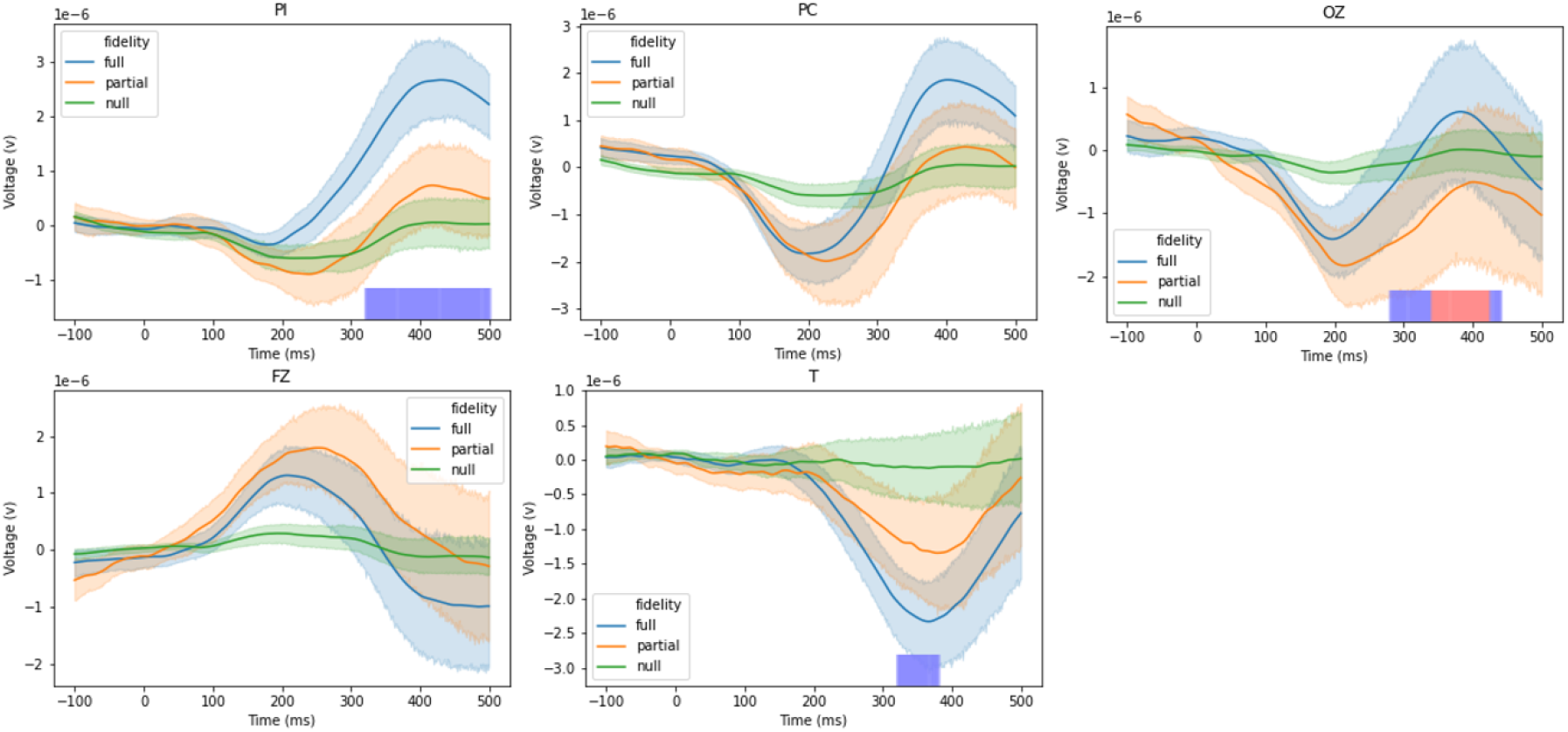
Average ERPs after performance-based adjustment and smoothing. These signals are aligned by the onset of the contour embedded frame, as represented by 0ms. Shadings indicate the 95% of confidence interval across all subjects. Different colors represent three contour fidelity conditions. The purple and pink squares lying on the X-axis mark the latency period when RM-ANOVA detected a significant effect from fidelity conditions (full vs partial) on ERP amplitudes after being corrected by an interval of 20ms with a critical value of 0.05 (purple) and 0.01 (pink).

### 3.3. Conclusions

Previous studies have demonstrated the findings of P400 signals in parietal and occipital regions in response to consciousness and task-driven functions, such as mental efforts in working memory (see details in the Discussion section). The N300 in frontal and central regions has been recently reported in tasks related to object recognition and semantic object identification [Schendan & Kutas, 2007; Kumar et al., 2021; Chen et al., 2022]. However, to the best of our knowledge, this is the first study where temporal N300 is reported in the context of contour integration. We therefore propose this as a signature for grouping and integration processes involved in contour computation.

## 4. Discussion

In our experiments, we observed multiple ERP signatures that correlate with sensory and cognitive processes taking place during contour integration and identification. Here we discuss the previous research findings in relation to these signatures, and to interpret them with respect to our experiments so as to uncover the cortical activities in relation to our contour integration task. The frontal P200 has been proposed to reflect processes involved in selective attention and cognitive execution in working memory [Hillyard et al, 1973; Wongupparaj, 2018; Zhao et al., 2013]. As for the N200 and P400 components observed in the occipital cortex, major theories indicate that when visual stimuli enter awareness, they tend to elicit a more pronounced posterior negativity at around 200ms compared to stimuli that fail to enter awareness [review: Förster et al., 2020]. This discernible distinction has been referred to as “visual awareness negativity” (VAN) [Koivisto & Revonsuo, 2010]. Additionally, this VAN is associated with another positive signature after 300ms, which is also termed “late positivity” (LP) [Förster et al., 2020]. Comparable distinctive patterns were likewise detected within the posterior parietal cortex, where the N200 activity was only evoked by the target situated contralateral to the visual field (termed as N2pc), while the subsequent P400 activity occurred irrespective of the visual field. The N2pc has been widely interpreted as the manifestation of the mechanisms used to suppress distractors and select targets during top-down attentional processes [Eimer, 1996; Luck & Hillyard, 1994; Mazza et al., 2009], whereas the LP signatures in both parietal and occipital areas are still in debate [Förster et al., 2020]. Multiple hypotheses have been discussed for the potential functional roles of LP. For example, one hypothesis considers this signature as a follow-up of the prior negativity, or a reflection of subjective awareness and visual consciousness [Dehaene & Changeux, 2011; Dehaene, 2014]. A second hypothesis states that this LP encodes the initiation of response after stimulus perception [Verleger et al., 2005]. Another hypothesis proposes that LP encodes the task-relevant effort and associated memory processing [Polich, 2007; Rac-Lubashevsk & Kessler, 2019].

Our results support the view of task-related effort and memory processing as the magnitudes of LP in both occipital and posterior parietal cortex were dependent on contour fidelity, whereas both VAN and N2pc were not. We believe that the LP signals observed in our experiments most likely signify a process of object recognition, where the perceived contour is compared to the complete contour stored in memory. In addition, we observed a novel ERP component in the context of contour integration, the temporal N300, which was evident in both sides of temporal cortex and exhibited an amplitude that was contingent on the fidelity of the contour. In the time domain, this signature fell in the duration after the signatures related to visual awareness and attentional effects and before the LP. It suggests that this signature was induced by integrating the visual elements into a contour.

Traditional views on the hierarchical neural structures in assembling local features into a global contour emphasized roles of the primary visual cortex. Multiple previous studies have reported that neural responses from V1 in macaque monkeys were correlated with contour integration [Roelfsema et al., 1998; Bauer & Heinze, 2002; Roelfsema et al., 2004; Li et al., 2006]. However, as a mid-level visual processing linking lower level feature binding to higher level object recognition, contour integration has been suggested to be associated with multiple mechanisms such as feature saliency, attention, and memory, etc., through interplays between different cortical areas [Chen et al., 2014; Shpaner et al., 2013; Silverstein et al., 2015]. On the other hand, the problem of contour integration becomes more complex in the real world as objects are perceived against a background, which requires coordination between contour integration and background segregation. Early studies using human psychophysics demonstrated that stimuli that conspicuously stand out against their background can capture attention and thus can be identified effortlessly, without the need to search through every element in the display (bottom-up processing) [Egeth, Jonides, & Wall, 1972; Jonides & Yantis, 1988; Theeuwes, 1994]. In comparison, stimuli that lack distinct saliency necessitate intentional direction of attention and a step-by-step examination of elements in the display before they can be recognized as search targets (top-down processing) [Wolf & Horowitz, 2004]. These attentional deployments have been widely reported to be correlated with activities in frontal, posterior regions, and their connections [Szczepanski et al., 2010; Li et al., 2010; Buschman & Miller, 2007; Constantinidis & Steinmetz, 2001; Ipata et al., 2006; Thomas & Paré, 2007; Knight, 1997; van Schouwenburg et al., 2010; van Schouwenburg et al., 2015; Szczepanski et al., 2014]. In the context of contour integration, both attentional processes have been suggested to play important roles to achieve effective contour-integration [Li et al., 2006; Li et al., 2008; Chen et al., 2014]. However, the interplay between different cortical areas to deploy these processes during the integration of contour is still largely unknown.

Our results show that the lateral frontal and posterior parietal regions are projected in an interhemispheric manner, with lateralization being dependent on the visual field of the target. Temporally following these activities, we observed a reduction in correlations between median frontal and superior parietal activities, the extent to which was contingent on the event order. This phenomenon can be explained by changing attentional effort with respect to uncertainty. When the contour is observed in the earliest possible frame, our perceptual system may maintain a certain level of top-down attention to continue monitoring, as it could be a false alarm. However, when the contour is observed in the last possible frame, confidence reaches the highest level, and thus we are likely to show the greatest reduction in top-down attention. At the same time, connections between lateral frontal and posterior parietal cortexes in one or the other side of the hemisphere, occipital and posterior parietal cortexes, as well as occipital and prefrontal cortexes were observed. Taking all these findings together, we suggest that bottom-up attentional effort directed by feature saliency is represented in an interhemispheric lateral frontal-posterior parietal pathway, whereas a posterior parietal-medial frontal pathway is responsible for encoding top-down fundamental attentional control.

